# Spatial and Amplitude Dynamics of Neurostimulation: Insights from the Acute Intrahippocampal Kainate Seizure Mouse Model

**DOI:** 10.1101/2023.03.07.531440

**Authors:** Thomas J. Foutz, Nicholas Rensing, Lirong Han, Dominique M. Durand, Michael Wong

**Affiliations:** Department of Neurology, Washington University School of Medicine, St. Louis, MO; Department of Biomedical Engineering, Case Western Reserve University, Cleveland, OH

**Keywords:** neuromodulation, seizures, epilepsy, status epilepticus, brain stimulation

## Abstract

**Objective:** Neurostimulation is an emerging treatment for patients with medically refractory epilepsy, which is used to suppress, prevent, and terminate seizure activity. Unfortunately, after implantation and despite best clinical practice, most patients continue to have persistent seizures even after years of empirical optimization. The objective of this study is to determine optimal spatial and amplitude properties of neurostimulation in inhibiting epileptiform activity in an acute hippocampal seizure model.

**Methods:** We performed high-throughput testing of high-frequency focal brain stimulation in the acute intrahippocampal kainic acid mouse model of temporal lobe epilepsy. We evaluated combinations of six anatomic targets and three stimulus amplitudes.

**Results:** We found that the spike-suppressive effects of high-frequency neurostimulation are highly dependent on the stimulation amplitude and location, with higher amplitude stimulation being significantly more effective. Epileptiform spiking activity was significantly reduced with ipsilateral 250 μA stimulation of the CA1 and CA3 hippocampal regions with 21.5% and 22.2% reductions, respectively. In contrast, we found that spiking frequency and amplitude significantly increased with stimulation of the ventral hippocampal commissure. We further found spatial differences with broader effects from CA1 versus CA3 stimulation.

**Significance:** These findings demonstrate that the effects of therapeutic neurostimulation in an acute hippocampal seizure model are highly dependent on the location of stimulation and stimulus amplitude. We provide a platform to optimize the anti-seizure effects of neurostimulation, and demonstrate that an exploration of the large electrical parameter and location space can improve current modalities for treating epilepsy.

**Key Points:**

- Evaluated spatial and temporal parameters of neurostimulation in a mouse model of acute seizures
- Brief bursts of high-frequency (100 Hz) stimulation effectively interrupted epileptiform activity.
- The suppressive effect was highly dependent on stimulation amplitude and was maximal at the ipsilateral CA1 and CA3 regions.
- Pro-excitatory effects were identified with high-amplitude high-frequency stimulation at the ventral hippocampal commissure and contralateral CA1.

## Introduction

Epilepsy affects 1-2% of all people, among whom about 30%^1^ have drug-resistant epilepsy (DRE) characterized by intractable seizures despite appropriate pharmacologic treatment^2^. The most common form of focal DRE is mesial temporal lobe epilepsy (MTLE)^3^, in which surgical removal can often treat the epileptic brain tissue to eliminate seizures in up to 70% of patients^4^. Neurostimulation therapy provides an alternative therapeutic option for those that are not candidates for surgical resection^5^, and can be less invasive, reversible, and customizable after implantation. The FDA-approved neurostimulation therapies which have demonstrated efficacy in treating refractory epilepsy are Vagus Nerve Stimulation (VNS)^6^, Deep Brain Stimulation (DBS)^7,8^, and Responsive Neural Stimulation (RNS)^9,10^. Unfortunately, several studies have shown that neurostimulation typically provides only modest reductions in seizure burden, rarely provides seizure freedom^11^ and requires numerous sessions over several years to attempt to optimize stimulation parameters. There are significant limitations in our knowledge about the underlying mechanisms and factors affecting the therapeutic effect of neurostimulation that limit our use of this therapy.

Optimization of neurostimulation for epilepsy involves simultaneously minimizing the disruption of normal neuronal activity, maximizing the disruption of abnormal activity (seizures), and avoiding the generation of new non-physiologic activity. This has been challenging due to non-linear responses to parameter changes and a poor understanding of the therapeutic mechanisms. Nevertheless, there is mounting evidence that neurostimulation acts by direct inhibition of neuronal activity^12^, activation of presynaptic inhibitory inputs^13^, and activation of efferent projections^14^. High-frequency stimulation (HFS, ~100-165 Hz) may also have thermal effects which can induce a transient suppression of neurons^15^. While HFS has been used more extensively^16^, low-frequency stimulation (LFS, ~1-10Hz) has significant promise for treating epilepsy^17,18^. Despite these previous studies, the optimal methodology for adjusting the stimulation frequency and target has not yet been found.

Current available brain stimulation platforms (DBS, RNS) stimulate along only one or two trajectories and are therefore highly spatially limited to where they directly apply electrical current. Stimulation generates an electric field around the active contacts, which decays in a roughly spherical distribution^19^. The size and shape of modulated neural activities depends on stimulation parameters^20,21^. Two spatial strategies for neurostimulation have emerged over the last several decades: direct stimulation of the suspected seizure-onset zone, and stimulation at a remote site that is involved in the seizure-onset network. RNS is typically used to directly target the suspected seizure-onset zone (e.g., hippocampal or cortical), and DBS to stimulate network hub locations (e.g., thalamic nuclei); however, such a distinction is increasingly become less well defined^22,23^. In the case of mesial temporal lobe epilepsy (MLTE), there are several attractive targets that have been explored, including the hippocampus^5,24–26^, the hippocampal commissures, and the medial septum. The hippocampal formation has a well-characterized functional network, where in general, CA3 neurons project to the CA1 region, and output via the subiculum. Anti-seizure effects have been found with stimulation at each of these regions^27,28,28–32^ as well as at hippocampal white-matter commissures^33,34,18^. Therapeutic effects have also been seen with stimulation of extra-hippocampal structures, including the anterior nucleus of the thalamus^7^ and medial septum^35,36^. Only limited *in vitro* studies^37^ have evaluated head-to-head comparisons of the CA1, CA3, and subiculum*; in vivo* studies are needed to more fully understand the spatial dependence of neurostimulation in the intact hippocampal network. In addition, the neuromodulatory effect of neurostimulation is likely to be highly dependent on the selected parameter settings^38–40^.

Because of the time required to evaluate effects on seizures, clinicians rarely can test even a small proportion of the vast stimulation parameter space (defined by the variation of active electrode location, current amplitude, pulse width, frequency, and burst duration) before settling on a long-term setting. By comparison, animal models of epilepsy address many challenges inherent to studying neurostimulation in humans, including heterogeneity in etiology and seizure characteristics, and allow high throughput testing of stimulation parameters. In this study, we utilized the acute intrahippocampal kainate seizure model to identify distinct spatial differences in (1) where spikes arise, (2) where stimulation is applied, and (3) how stimulation amplitude modulates the therapeutic effects of direct hippocampal stimulation.

## Materials & Methods

### Acute Intrahippocampal Kainate Mouse Model of Focal Cortical Seizures

Adult male and female mice with mixed C57Bl/6 and SV129 genetic backgrounds were used in this study (n=19, four females). Animals were anesthetized for the duration of the experiment using isoflurane and regularly monitored to ensure appropriate anesthesia. Burr holes 0.9 mm in diameter were then drilled through the skull overlying each target brain location. A pulled glass pipette was slowly lowered via stereotactic frame into the right dentate gyrus region of the hippocampus (Bregma AP: −2, ML: 1.5, DV: −1.8 mm). Seizure activity was induced by three small bolus injections of KA (total 75 nL x 20 mM) using a nanoliter injector (NanoJect III, *Drummond Sci*), injected over 10 minutes. After the completion of stimulation trials, animals were euthanized and in a subset of cases (n=5), histological verification of electrode placement was done from the tissue artifact seen on cresyl violet stained 50 μm slices. The Washington University School of Medicine Institutional Animal Care and Use Committee reviewed and approved the experimental protocols.

### Electrode Placement and Electroencephalography

After KA injection, four skull-based stainless screw electrodes were implanted over the frontal and parietal bones. Two additional screw electrodes were placed over the interparietal bone for ground and reference. Then, six custom 0.127 mm diameter platinum-iridium wire electrodes, sheathed in fused silica, were stereotactically lowered through their respective burr hole into target structures for both stimulation and recording (Figure 1A-B), with the following Bregma Coordinates (AP; ML; DV): Bilateral CA1/Dentate Gyrus (CA1: −2.5; ±1.7; −1.6); Ipsilateral Subiculum (SUB: −2.5; −0.5; −1.7) and CA3 (CA3: −2.5; −2.8; −2.6); Ventral Hippocampal Commissure (VHC: −0.7; +0.1; −2.2), and Medial Septum that had an angulated trajectory (MS: AP +0.6; ML +1 at 15°; −3.5 mm from the pial surface). Depth electrodes were secured with dental acrylic and cyanoacrylate. Electroencephalographic recordings (EEG) and electrical stimulation trials were then performed via a 16-channel headstage (M4016, *Intan Tech*) connected to a stim/recorder controller (M4200, *Intan Tech*) sampled at 25 – 30 kHz. Discrete seizures were observed in the first 30 minutes after injection, which evolved into nearly continuous epileptiform activity characterized by high-amplitude spiking overlying fast activity that lasted several hours, consistent with status epilepticus (Figure 1C). Acute neurostimulation was evaluated in this state of continuous spiking activity.

**Figure 1.**
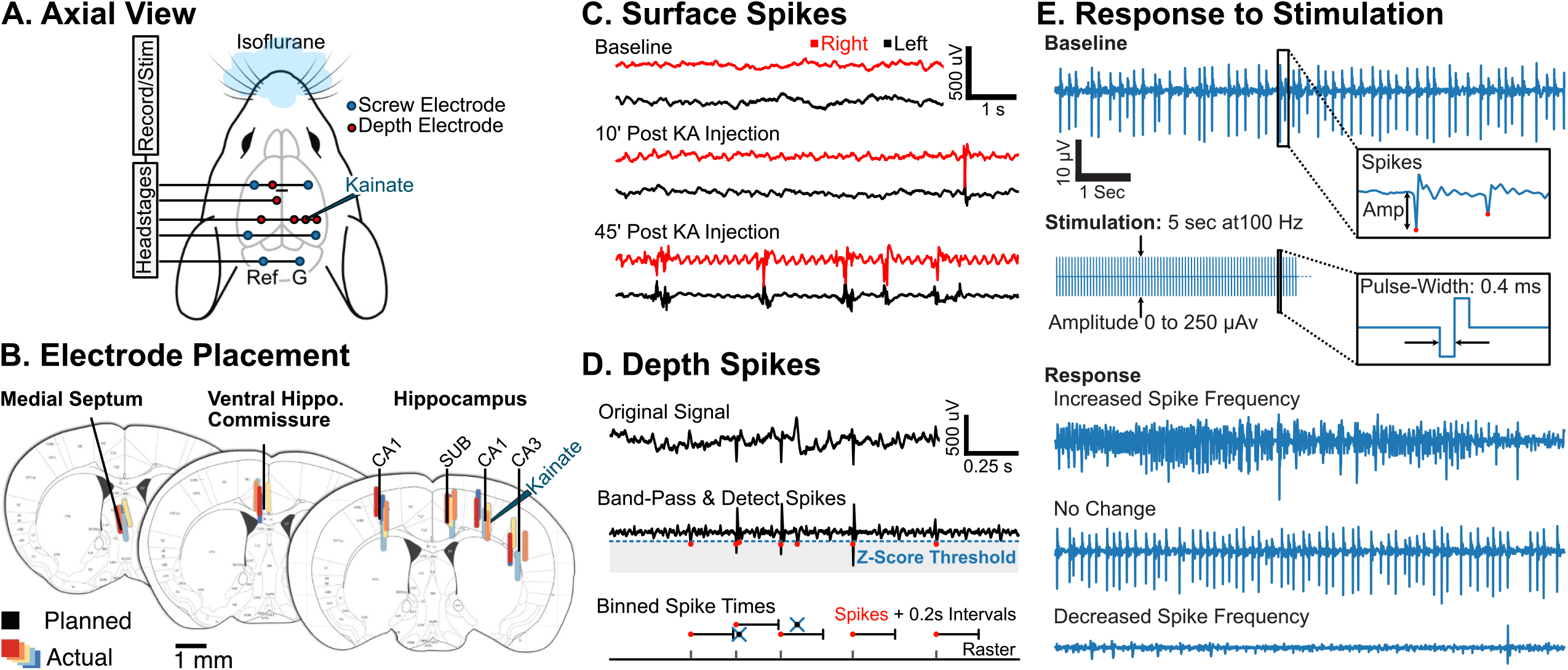
Electrode Implantation and Intrahippocampal Kainate Induced Spiking. **A.** Surface electrode implants, axial view. While under isoflurane anesthesia, six stainless-steel screw electrodes were implanted in the frontal-parietal and interparietal skull bones. **B.** Depth electrode implants, coronal views. KA was injected into the right-sided CA1/DG region. Then, six intracerebral depth electrodes were placed in the bilateral CA1, ipsilateral subiculum, CA3, ventral hippocampal commissure, and medial septum. Atlas images adapted from the original^50^. Colored rectangles represent approximate location of active electrode based on histolologic evaluation (n=5). **C.** KA-induced spiking activity evolution is seen on surface recordings. Bilateral surface EEG recordings (frontal-parietal; right and left sides) were performed during ictogenesis post-implantation. Over time, high-amplitude spike and poly-spike epileptiform discharges develop. **D.** Spike detection performed on depth recordings. After removing the common mode activity (top), the signal was band-pass filtered, and a modified z-score was estimated (middle). The threshold for spike detection was empirically set (z>5) to provide adequate spike detection while limiting false detection of non-epileptiform activity. Spikes occurring within 0.2 sec of the last detection were removed (bottom). **E.** Different effects of acute high-frequency stimulation were seen. *Baseline:* After KA infusion, spiking at the ipsilateral CA3. The inset demonstrates two detected spikes with amplitude (Amp). *Stimulation:*High-frequency (100 Hz) stimulation was applied for 5 seconds during each trial, with a range of amplitudes from 0 (control), to a maximum of 250 μA. The EEG recording is overwhelmed by electrical switching artifacts and was not used in the analysis. *Response:* Example traces demonstrating spike frequency changes after stimulation: increased spike frequency, no appreciable change, and decreased spike frequency.

### Application of Different High-Frequency Stimulation Protocols

A range of stimulation amplitudes was evaluated in each target structure across multiple repeat blocks of trials. Before starting stimulation, the electrode impedances were verified to be within acceptable limits (500 Ω – 28 kΩ) to permit generating the planned range of stimulus amplitudes (0 – 250 μA). Stimulus charge per phase was limited to safe levels that have previously been established (<30 μC/cm^2^ per phase)^41^ to ensure that the neuromodulatory effect is due to reversible activity rather than irreversible tissue damage; no tissue charring or other visible evidence of electrical tissue injury was seen on histology. Then, repeat stimulation trials were performed, drawn from a range of stimulation amplitudes (0, 50, 100, 250 μA) and electrode locations (MS; VHC; Bilateral CA1; Ipsilateral SUB & CA3). The parameter sequence was randomly shuffled for each block of trials. Each trial started with a 90-second epoch of recording baseline activity. Then, current-controlled stimulation was administered by the headstage at a frequency of 100 Hz for 5 seconds with a 400 μsec pulse-width, cathodic-first charge-balanced waveform. Trials with 0 μA amplitude acted as controls. Finally, a 90-second epoch of recording was performed to capture the post-stimulation activity. The time between stimulations was 3.2 minutes or greater (*M* = 205 s, *SD* = 174). Furthermore, trials were randomized to minimize carry-over effects from prior trials.

### Signal Processing and Spike Detection

EEG recordings were reviewed during the recording to verify appropriate signal quality and anesthesia. A total of 1433 trials were acquired, with a mean of 75.1 trials per mouse (*SD* = 57.5) and including 132 no-stimulation control trials. Recordings were band-pass filtered (20 – 500 Hz), and the common mode signal was removed from surface and depth recording datasets. For spike detection, a modified z-score^42^ was calculated from the baseline activity to allow for consistent spike detection before and after stimulation, defined as:

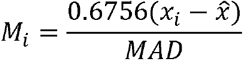

where *x_i_* is the signal at index *i*, and 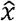 is the median of the baseline signal. MAD is the median absolute deviation, defined as:

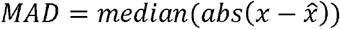

Where *x* is the baseline signal, and 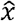 is its median value. As has been done in similar electrophysiologic studies for spike detection, negative amplitude spikes were detected when the modified z-score of amplitude (*M_i_*) had a magnitude of five or greater^43,44^. Depth electrodes were used for spike counts and stimulation, whereas surface EEG screw electrodes were used as reference to reduce common mode signal and EKG artifacts. The time-to-100^th^ spike (TTH) was defined as the time required for 100 spikes to be detected. Only trials with 100 spikes in the baseline period were included (n=1233). In a few cases (n=58), less than 100 spikes were detected in the post-stimulation period; in these cases, the TTH was set to the upper limit (90 sec). Outcomes were classified as (a) *Improved* if the TTH was increased by 25% or more in the post-stimulation epoch compared to baseline, (b) *Worsened* if the TTH decreased by 25% or more, else (c) *No Change*.

Spike data was further analyzed to determine individual spike characteristics and changes in spike frequency over time. Spike amplitude was calculated as the maximum negative amplitude compared to the baseline signal 50-25 milliseconds before the peak detection^45^. Since spike detection was based on pre-stimulation baseline, increased post-stimulation spike amplitude alone should not affect the detected spike frequency. The baseline spike frequency was defined by the mean spike count per second over the final 20 seconds pre-stimulation, whereas the post-stimulation spike frequency was defined over the initial 20 seconds post-stimulation. When comparing post-stimulation spike frequency to baseline, to permit evaluation of both increased and decreased changes in spike frequency, trials (n=896) were required to have a mean of at least two spike/s. Spike-frequency over time in the post-stimulation epoch was further evaluated using spike-time density plots for each stimulus amplitude and location combination.

Spikes often had a field extending into other electrodes or had a poly-spike morphology, which could lead to duplicate counting. Therefore, the spike count was performed in each epoch (baseline and post-stimulation) after detecting the spike onset and discarding any detections in the subsequent 200 milliseconds. This counting method allowed for combined-spike rate detection of up to five spike periods per second, which was used for quantification of the TTH, spike rates, and onsets (Figures 3–6).

**Figure 2.**
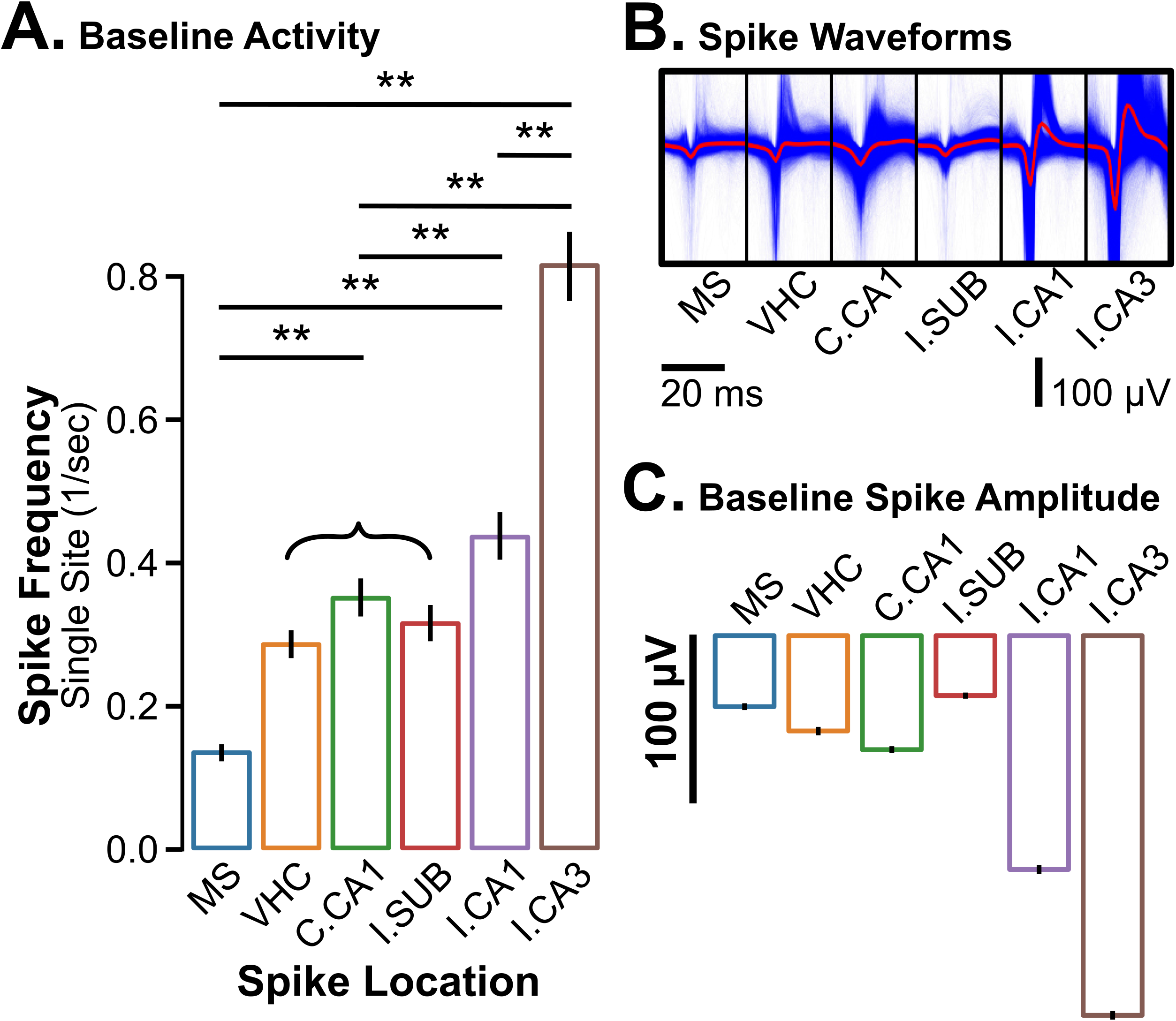
Spike Characteristics during baseline recording. A total of 413,323 spikes were detected and analyzed across all trials. **A.** Spike frequencies were compared at each location. Significant differences were found between spike frequencies at most sites (* *p < .001*); however, no differences were found between VHC, contralateral CA1, and ipsilateral subiculum, so these are shown combined. **B.** Single and averaged waveform traces are shown for each depth location. The highest spike amplitudes are seen at the ipsilateral CA3 region. **C.** Mean baseline spike amplitudes. Amplitudes at each site were significantly different (*p < .001*). MS: Medial Septum, VHC: Ventral Hippocampal Commissure, SUB: subiculum, C.: Contralateral CA1, I.: Ipsilateral. Error bars represent a 95% confidence interval.

**Figure 3.**
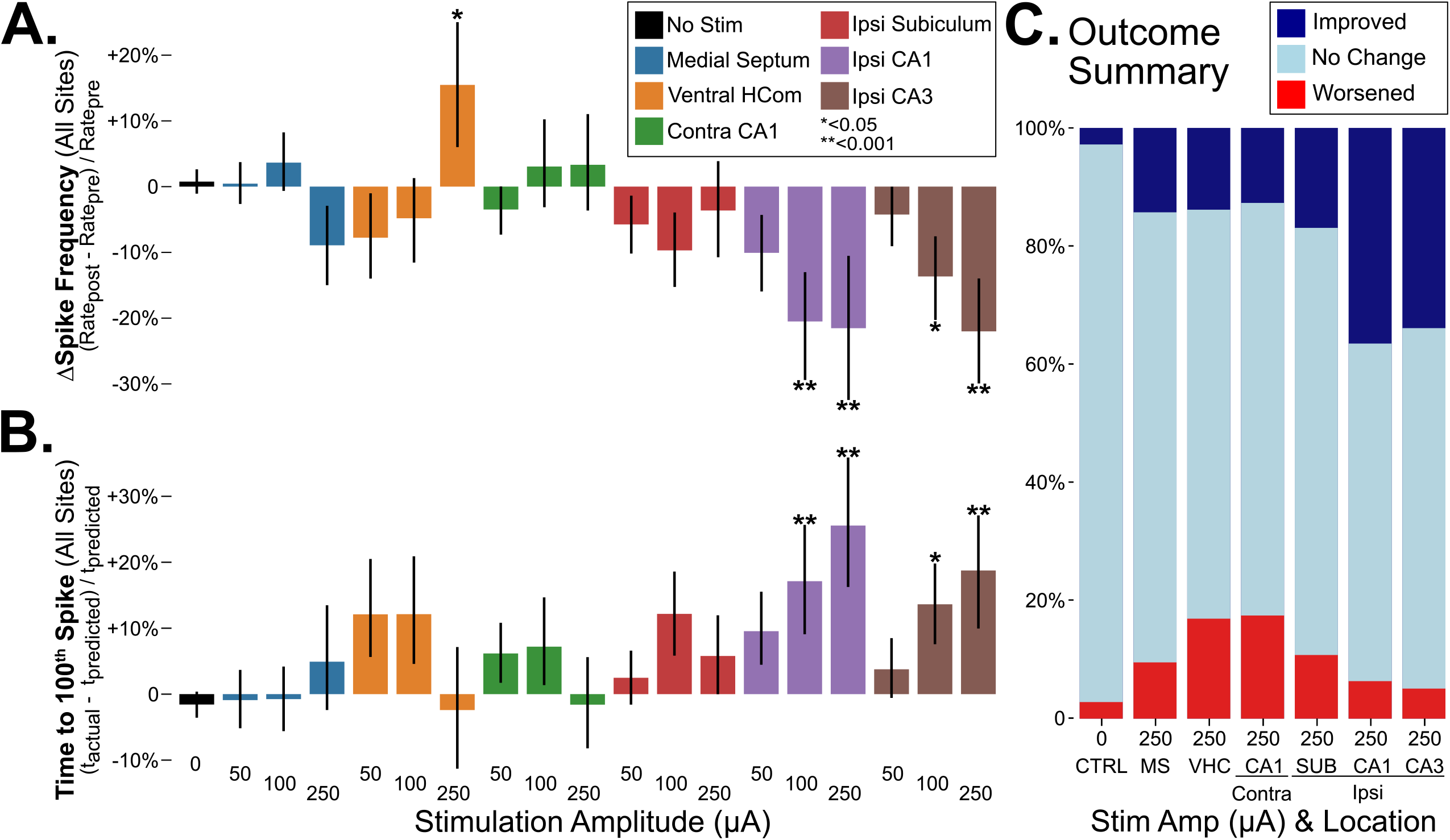
Acute High-Frequency Stimulation Effects on Epileptiform Activity Between Different Targets and Amplitudes. **A.** Percentage change in combined-spike frequency, compared to no-stimulation control. (* p<0.05; ** p < 0.01). **B.** Percentage change in the TTH, compared to control. **C**. Categorized outcome to maximal amplitude (250 μA) stimulation at each electrode location. The response was categorized as “Improved” if the TTH spike was greater than or equal to 125% of baseline; as “Worsened” if less than or equal to 75% of baseline; otherwise, as “No Change.” CTRL: Control/No-Stim. MS: Medial Septum, HCom: Hippocampal Commissure, VHC: Ventral HCom, SUB: subiculum, Contra: Contralateral, Ipsi: Ipsilateral. Error bars represent a 95% confidence interval.

**Figure 4.**
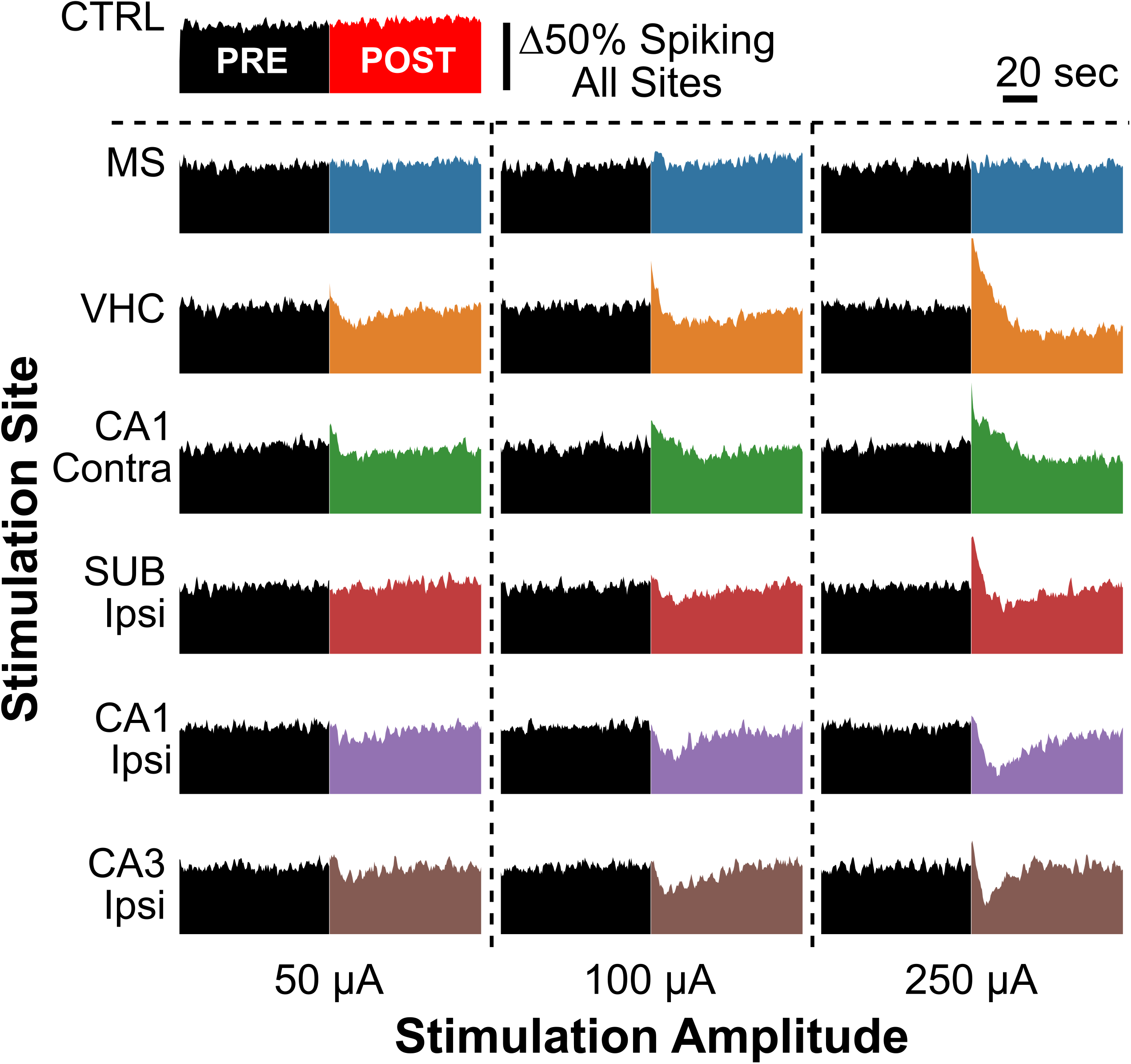
Spike-Time Density Before and After Stimulation Between Different Targets and Amplitudes. Spikes were binned into 1-second epochs and averaged across all trials for each combination of stimulus and location. The height of the density plots corresponds to a relative change compared to the mean baseline spike frequency. Stimulation locations are listed on the left, and stimulation amplitudes are on the bottom. The black density plot represents the combined-spiking activity before stimulation, whereas the color-coded density plot represents the combined-spiking activity post-stimulation. CTRL: Control/No Stim. MS: Medial Septum, VHC: Ventral Hippocampal Commissure, SUB: subiculum, Contra: Contralateral, Ipsi: Ipsilateral.

**Figure 5.**
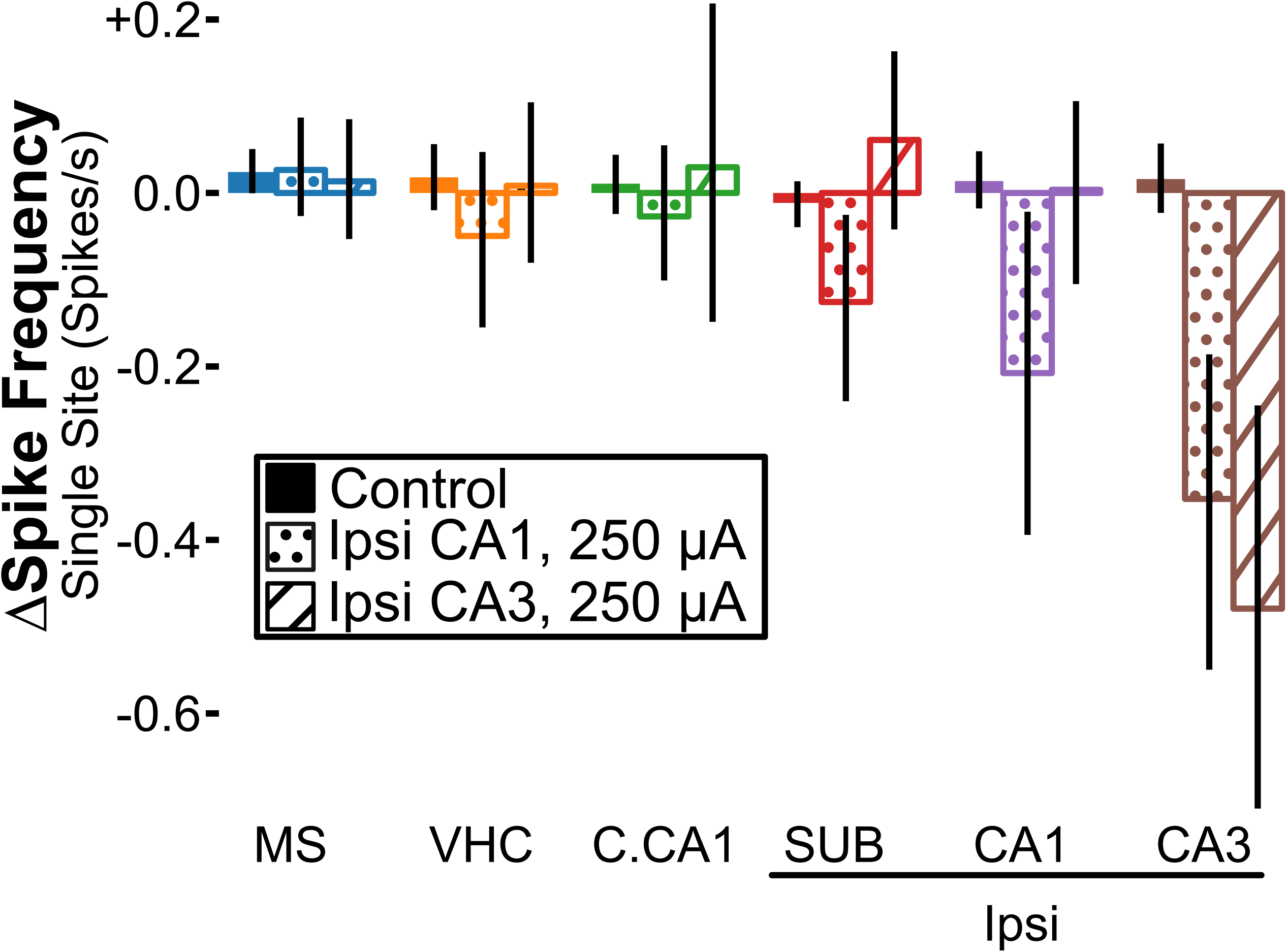
Spatial Response to Stimulation for Each Site using Effective Stimulation Parameters. The spike-suppression effect was evaluated at each recording site in response to the two stimulation parameter sets that consistently showed a significant reduction in spiking (Ipsilateral CA1 & CA3, 250 μA stimulation amplitude). Results were compared to the control. Error bars represent 95% confidence intervals. MS: Medial Septum, VHC: Ventral Hippocampal Commissure, SUB: subiculum, C.CA1: Contralateral CA1, Ipsi: Ipsilateral.

**Figure 6.**
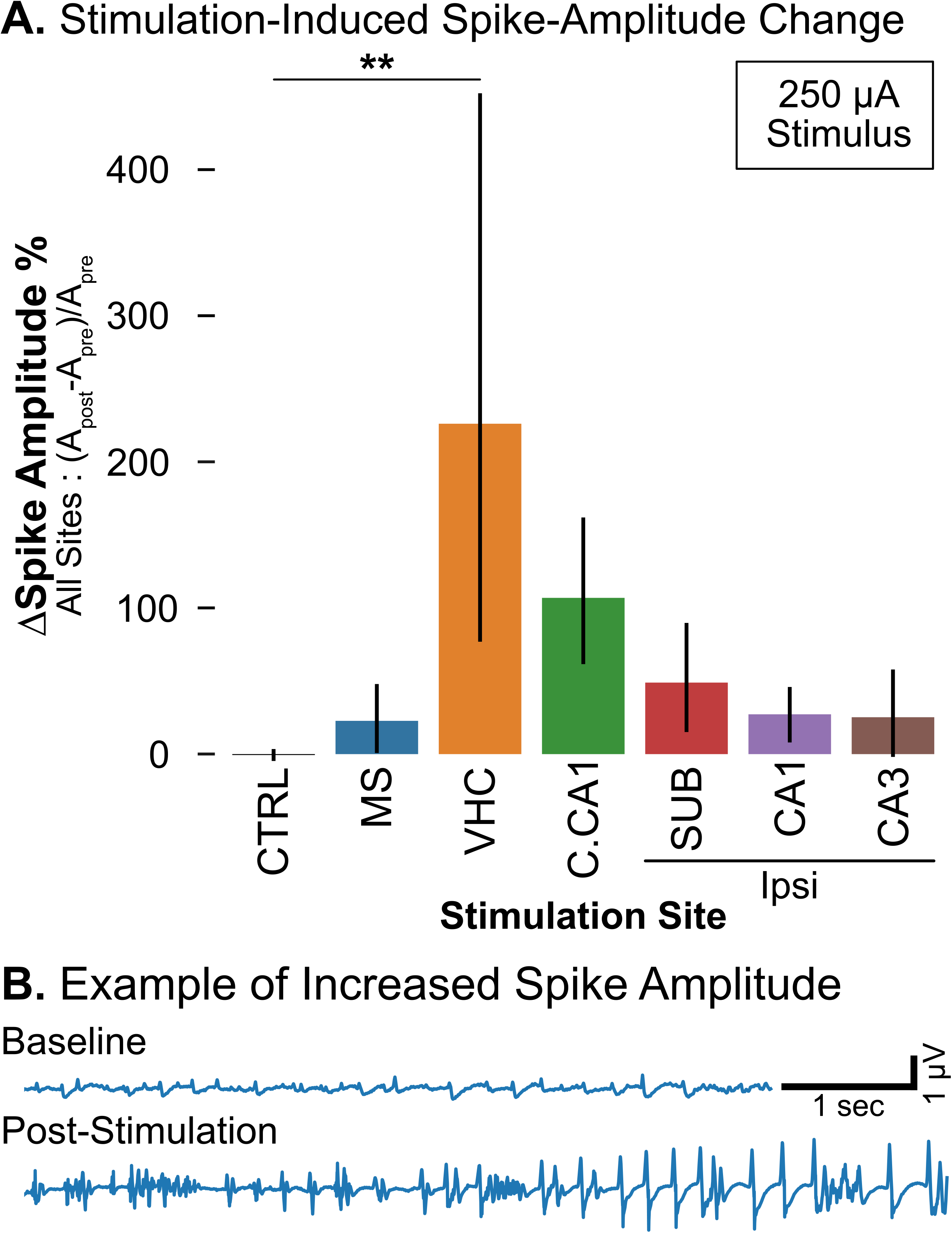
Spike Amplitude Changes after Maximal Stimulation: Comparison Between Different Stimulation Targets. **A.** Post-stimulation spike amplitudes were averaged across all recording sites and normalized to baseline. Maximal Stimulation (250 μA) was applied at each location. Mean spike amplitudes across all sites are increased for stimulation at VHC (** p<.01). CTRL: Control/No Stim. MS: Medial Septum, VHC: Ventral Hippocampal Commissure, SUB: subiculum, C.CA1: Contralateral CA1, Ipsi: Ipsilateral. Error bars represent a 95% confidence interval. **B.** Example recording before and after HFS 100 μA stimulation at the contralateral CA1. Note the increased spike frequency and amplitude post-stimulation.

### Statistical Analysis

Data were analyzed as a two-way ANOVA linear model using the *anova_lm* function in the python *statsmodels* toolbox^46^. Each trial was treated as an independent trial compared to its baseline. If ANOVA detected a significant difference, the *multicomp.pairwise_tukeyhsd* function in *statsmodels* was used to calculate pairwise comparisons with Tukey honestly significant difference (HSD) confidence intervals, accounting for multiple comparisons. For both tests, the significance level *alpha* was set to 0.05, and a significant difference was defined as *p* less than or equal to 0.05.

## Results

### Baseline Activity and Spike Characteristics

During the pre-stimulation baseline, the average combined-spike frequency across all animals and trials was 2.4 spike/s (*SD = 1.0*). Across all animals, the mean baseline spike frequency started at 2.2 spike/s, increasing to a peak of 3.0 spike/s over the first 1.8 hours, and then gradually dropping below 2 spike/s at 6 hours. Spikes were most often first detected at the ipsilateral CA3 (*M = 0.82 spike/s*), followed by ipsilateral CA1 (*M = 0.44*), and least frequent at Medial Septum (*M = 0.13*), as shown in Figure 2A. All spike waveform tracings (blue) are plotted in Figure 2B, overlaid with the average spike tracing (red). The mean spike amplitude, defined as the peak negative amplitude of the spike compared to the baseline (Figure 1E inset), was - 130 μV (*SD = 160*) across all animals and trials. The spike amplitude varied significantly between sites (Figure 2C) and was found to be most prominent at the ipsilateral CA3 (*M = −227 μV, SD = 181*) and least prominent at the ipsilateral subiculum (*M = −35.8 μV, SD = 43.5*).

### Spike Frequency and Time to 100^th^ Spike

The spike detection frequency varied significantly between stimulation sites. After excluding trials with low spike frequencies (i.e., <2 spikes/s), the included trials (n=896) had a baseline combined-spike frequency (pooled from all depth electrodes) of 3.0 spike/s (*SD=0.59*). After the no-stimulation sham period in trials randomized to 0 μA stimulation, the combined-spike frequency remained essentially unchanged at 3.0 spike/s (*SD=0.56*), with an estimated poststimulation percentage change of +0.98%. Combined-spike frequencies were similarly evaluated across all eighteen stimulation amplitudes and location combinations. After comparing results using ANOVA and Tukey tests, only five combinations demonstrated significantly different percentage changes compared to the no-stimulation control group (Figure 3A). VHC 250 μA stimulation demonstrated a worsening spike-frequency *increase* of +15.3% (*p < .05*). In contrast, stimulation with either 100 or 250 μA, at CA1 or CA3, demonstrated improvement with significant spike-frequency *decreases* between −13.0% to −22.2% (*p < .05* for 100 μA at CA3, otherwise *p < .01*).

Similarly, the TTH was compared across all stimulation combinations. The percentage change was determined by comparing each trial’s post-stimulation TTH to their respective predicted TTH, estimated by 100 spikes divided by the baseline mean spike frequency. For the control group with no stimulation, the predicted TTH was 42.5 sec (*SD=16.0*). After the no-stimulation sham period, the actual TTH was 41.3 sec (SD=14.3) with a non-significant estimated percentage change of −1.6% (*SD=9.5*). After comparing results using ANOVA and Tukey tests, four combinations demonstrated significant TTH percentage changes from the no-stimulation control group. These were all combinations of 100 and 250 μA at locations ipsilateral CA1 and CA3, which demonstrated improvement with a significant *prolongation* of TTH between 13.6–25.6% (*p < .05* for 100 μA at CA3, otherwise *p < .01*), as presented in Figure 3B.

The response to 250 μA stimulus amplitude at each location was compared in Figure 3C. For the no-stimulation control case, 94.4% of trials demonstrated no significant change in spiking (*TTH_post_~TTH_pre_*), 2.8% with increased spiking (*TTH_post_≥ 1.25 * TTH_pre_*), and 2.8% with decreased spiking (*TTH_post_ < 0.75 * TTH_pre_*). Stimulation at the contralateral CA1 demonstrated the highest percentage of trials that had worsened TTH (*16.9%*), associated with increased trials that had improved TTH (*13.8%*). In contrast, stimulation at the ipsilateral CA1 demonstrated the highest percentage of trials that had improved TTH (*36.5%*), with a marginal increase in trials that had worsened TTH (*6.3%*). Ipsilateral CA3 stimulation demonstrated similar improvement in TTH.

### Spike Time Density

To better understand the temporal changes in spiking activity, spike frequencies were charted over time in a density plot (Figure 4). These plots demonstrate that the spike-suppressive effect of neurostimulation is time-limited, showing gradual recovery back to the baseline spike frequency. Qualitatively, most stimulation settings demonstrate an initial transient increase in spike frequency; only at a subset of sites (ipsilateral CA1, CA3) is an overall reduction in spike frequency appreciated. For the maximum amplitude stimulation at ipsilateral CA1 or subiculum, the spike frequency approached back to baseline around the end of the trial, whereas for ipsilateral CA3 the recovery was faster, by around 40 seconds. No appreciable change from baseline spiking frequency was seen with MS stimulation. For stimulation at the contralateral CA1 region or ventral hippocampal commissure, the spike frequency approached back to baseline by the end of the post-stimulation recording phase (90 seconds) for low-amplitude (50, 100 μA) stimulation, but for 250 μA stimulation the spike frequency remained slightly depressed at the end of the recording phase. However, on a review of the trials that immediately followed, there is no appreciable difference in spiking activity compared to control trials, nor is there any appreciable evolution in spike frequency over their baseline period.

### Spatial Distribution of Spike Suppression Depends on Stimulation Location

Though stimulation was applied similarly at each site, the spatial effects of stimulation was different (Figure 5). Stimulation at the ipsilateral CA1 with 250 μA stimulus amplitude generated significant reductions in spike frequency at multiple sites, including the ipsilateral subiculum, CA1, and CA3. However, when the same amplitude of stimulation was applied to the ipsilateral CA3, a significant reduction in spike frequency was seen only at a single site, the ipsilateral CA3. No significant changes were seen for control trials.

### Post-Stimulation Spiking Characteristics

Evaluation of post-stimulation spiking revealed that across all locations, there was a trend towards slightly higher amplitudes (Figure 6A), but only with stimulation at the ventral hippocampal commissure was this increased amplitude found to be statistically different from no-stim control (*M = 2.3, SD = 8.2, p < .01*). Out of all recordings (n=1433), transiently increased repetitive high-amplitude spiking (modified z-score > 10) was seen in 55 trials (3.8%), with or without bursting. This high-amplitude activity was seen most prominently with contralateral CA1 stimulation (n=28, 2.0%) versus ipsilateral CA3 stimulation (n=2, 0.1%). This activity was also seen with stimulation at ipsilateral CA1, subiculum, and VHC (n=11, 7 and 7 respectively). A typical example of this high-amplitude spiking activity is shown in Figure 6B.

## Discussion

In this study, we tested the efficacy of high-frequency neurostimulation on acute intrahippocampal kainate-induced epileptiform spiking activity. This protocol was consistent in generating reproducible epileptiform spiking activity lasting several hours, consistent with status epilepticus. We found that high-amplitude (250 μA) stimulation of the ipsilateral CA1 and CA3 demonstrated significant spike-suppressive effects, prolonging the TTH and reducing the poststimulation spike frequency without significant effects on spike amplitude. Post-stimulation spike density plots show that this effect is transient. These data support that high-amplitude, high-frequency stimulation at CA1 and CA3 has an anti-seizure effect. Conversely, stimulation of the VHC region induced a significant increase in spike frequency, shortened the TTH, and generated a significant increase in post-stimulation spike amplitude, suggesting the potential for worsening seizure activity. Such an increase in post-stimulation amplitude is akin to that seen in human subjects during electrical stimulation mapping examinations, often referred to as “after-discharges,” which in this context represents additional excitation of already seizing brain. Optimal stimulation would ideally minimize this type of excitation. Together, these results indicate that high-frequency neurostimulation is highly spatially dependent, which has critical translational applications for treating epilepsy, particularly for future clinical studies that seek to optimize the seizure-suppressive effects of brain stimulation in hippocampal or other deep brain regions.

Brain stimulation therapy has been approved for treating epilepsy in the US and Europe; however, the precise mechanisms by which it reduces seizures are unknown. This lack of understanding hampers the development of rational approaches to improving stimulation settings when the therapy is ineffective. Suboptimal responses may be due to inappropriate spatial (e.g., active electrode location), amplitude, and temporal (e.g., frequency) parameters. While DBS has been approved for targeting the anterior nucleus of the thalamus, it is often used off-label to target several other locations with some evidence for benefit^25^. However, an exhaustive search of all potential brain targets, electric fields, and stimulus parameters has not been done, nor is such a study likely due to the enormity of potential parameter combinations. For this reason, animal studies can provide a platform for much higher-throughput exploration of the parameter space than is feasible in human studies. Stimulation trials can even be performed in multiple animals with identical electrode array layouts, which would be difficult to replicate in humans. The additional electrode recordings sites permit better characterizing spatial effects, which would be challenging to replicate clinically. Therefore, this animal work aims to provide a better understanding of the stimulation parameter space in a manner that informs the rational development of future clinical studies.

Among the stimulation locations and amplitudes tested in this study, stimulation at the ipsilateral CA1 and CA3 demonstrated a desirable response with decreased spiking activity and no significant increase in post-stimulation spiking amplitude. While not statistically different, there was an apparent dose-response relationship between low (100 μA) and high (250 μA) stimulation with increasing amplitude generating increasing effects. Since KA is thought to act primarily on KA receptors with the highest density in the CA3 region^47^, the finding that CA3 stimulation of this region produces a favorable effect lasting several seconds supports that high-frequency stimulation probably can act locally in the stimulated region. This effect may be due to the direct inhibition of excitatory neurons in the target region^48^, activation of local inhibitory neurons^16^, thermal effects^15^, or extracellular effects like potassium accumulation^37^. Alternatively, CA3 and CA1 may be better targets than output regions like the subiculum since HFS may primarily modulate downstream rather than upstream areas. This study does support that electrical stimulation may terminate or suppress ongoing seizures and that these effects depend on the amplitude and location of stimulation. It is challenging to evaluate the electrophysiologic changes during the 5-second stimulation period due to significant stimulation artifacts and saturation of the recording amplifiers. It is also unknown if higher amplitudes would further improve the response, or if paradoxical worsening would be seen. Higher stimulation amplitudes also risk inducing irreversible tissue damage.

The detrimental effect of high-frequency stimulation at the “mirror” CA1 focus and ventral hippocampal commissure is of interest, with similar effects that suggest a common pathway. One potential explanation for this is that under the KA-induced state, KA brings ipsilateral CA3 neurons close to their activation threshold such that excitatory inputs from the contralateral hemisphere worsen epileptiform activity. This finding contrasts with prior work demonstrating the benefit of low-frequency stimulation at the VHC^33,49^, suggesting that these effects are frequencydependent. The medial septum has a large body of evidence for reducing the frequency of seizures in chronic epilepsy models. However, in this acute model, no significant effect was seen with medial septum stimulation. Further studies while awake are planned to see if this effect may be limited to the unanesthetized state.

In this study, we found that, while both ipsilateral CA1 and CA3 demonstrated significant spike-suppressive effects, CA3 stimulation’s effects were limited to CA3 activity, whereas CA1 stimulation had more broad effects on the subiculum, CA1, and CA3 activity. These findings have important translational implications for patients who do not respond to electrical stimulation, possibly because their electrode is not close enough to the epileptogenic zone or because there are excitable regions outside the electrode’s network of influence.

One limitation of this study is that animals were maintained under isoflurane anesthesia; however, despite this anesthetic, the pathologic signature consistently seen across animals has a viable clinical counterpart of status epilepticus, maintaining its relevance to human epilepsy. Despite this limitation, this approach’s benefit is that many stimulation settings can be tested in a high-throughput manner. This study was further limited to six spatial locations; future studies will include additional targets of interest (e.g., anterior thalamus, CA2) and other stimulation parameters (e.g., frequency, pulse width, and stimulation polarity). Low-frequency stimulation (1 Hz, 50-250 μA) was performed early on in a subset of animals (data not shown); however, the preliminary analysis demonstrated no significant change in the spike frequency, so this portion of the parameter space was not pursued. It remains to be seen if other stimulation frequencies are as or more effective for stimulation in these regions. Finally, this study was limited to acutely-induced continuous epileptiform activity (status epilepticus). Future studies in chronic epilepsy models are needed to confirm that similar principles apply to spontaneous, recurrent seizures.

## Conclusion

Overall, the present study provides evidence that high-frequency hippocampal stimulation of the CA1 and CA3 regions suppress KA-induced epileptiform activity and may be good candidates for targeting deep brain stimulation for temporal lobe epilepsy, where CA1 stimulation demonstrates broader spike-suppressive effects. Conversely, high-frequency VHC stimulation was associated with detrimental increases in spiking activity and spike amplitude, which may need to be avoided. These findings have translational applications for designing clinical studies to evaluate spatial differences in response to high-frequency stimulation in epilepsy.

## Acknowledgements

This work was funded in part by the American Epilepsy Society Research & Training Fellowship for clinicians, an Institutional Postdoctoral T32 Training Grant provided by the National Institute of General Medical Sciences, NIH P50HD103525 to the Washington University Intellectual and Developmental Disability Research Center, and institutional support from Washington University in St. Louis.

## Author Contribution

T.F.: Conceptualization, Methodology, Validation, Formal Analysis, Investigation, Writing – Original Draft, Writing – Review & Editing; N.R.: Methodology, Investigation; L.H.: Investigation; D.D.: Conceptualization, Writing – Review & Editing; M.W.: Conceptualization, Writing – Review & Editing, Supervision, Project Administration.

## Conflict of Interest/Ethical Publication Statement

None of the authors has any conflict of interest to disclose. We confirm that we have read the journal’s position on issues involved in ethical publication and affirm that this report is consistent with those guidelines.

## References

1. Zack MM, Kobau R. National and state estimates of the numbers of adults and children with active epilepsy - United States, 2015. MMWR Morb Mortal Wkly Rep. 2017; 66(31):821–5.

2. Chen Z, Brodie MJ, Liew D, Kwan P. Treatment Outcomes in Patients With Newly Diagnosed Epilepsy Treated With Established and New Antiepileptic Drugs: A 30-Year Longitudinal Cohort Study. JAMA Neurol. 2018; 75(3):279–86.

3. Asadi-Pooya AA, Stewart GR, Abrams DJ, Sharan A. Prevalence and incidence of drug-resistant mesial temporal lobe epilepsy in the United States. World Neurosurg. 2017; 99:662–6.

4. Al-Kaylani M, Konrad P, Lazenby B, Blumenkopf B, Abou-Khalil B. Seizure freedom off antiepileptic drugs after temporal lobe epilepsy surgery. Seizure. 2007; 16(2):95–8.

5. Vonck K, Boon P, Achten E, De Reuck J, Caemaert J. Long-term amygdalohippocampal stimulation for refractory temporal lobe epilepsy. Ann Neurol. 2002; 52(5):556–65.

6. Morris GL, Mueller WM. Long-term treatment with vagus nerve stimulation in patients with refractory epilepsy. Neurology. 1999; 53(8):1731–1731.

7. Fisher R, Salanova V, Witt T, Worth R, Henry T, Gross R, et al. Electrical stimulation of the anterior nucleus of thalamus for treatment of refractory epilepsy. Epilepsia. 2010; 51(5):899–908.

8. Salanova V, Witt T, Worth R, Henry TR, Gross RE, Nazzaro JM, et al. Long-term efficacy and safety of thalamic stimulation for drug-resistant partial epilepsy. Neurology. 2015; 84(10):1017–25.

9. Morrell MJ, RNS System in Epilepsy Study Group. Responsive cortical stimulation for the treatment of medically intractable partial epilepsy. Neurology. 2011; 77(13):1295–304.

10. Heck CN, King-Stephens D, Massey AD, Nair DR, Jobst BC, Barkley GL, et al. Two-year seizure reduction in adults with medically intractable partial onset epilepsy treated with responsive neurostimulation: Final results of the RNS System Pivotal trial. Epilepsia. 2014; 55(3):432–41.

11. Touma L, Dansereau B, Chan AY, Jetté N, Kwon C-S, Braun KPJ, et al. Neurostimulation in people with drug-resistant epilepsy: Systematic review and meta-analysis from the ILAE Surgical Therapies Commission. Epilepsia. 2022; 63(6):1314–29.

12. Filali M, Hutchison WD, Palter VN, Lozano AM, Dostrovsky JO. Stimulation-induced inhibition of neuronal firing in human subthalamic nucleus. Exp Brain Res. 2004; 156(3):274–81.

13. Milosevic L, Kalia SK, Hodaie M, Lozano AM, Popovic MR, Hutchison WD, et al. A theoretical framework for the site-specific and frequency-dependent neuronal effects of deep brain stimulation. Brain Stimul. 2021; 14(4):807–21.

14. Hashimoto T, Elder CM, Okun MS, Patrick SK, Vitek JL. Stimulation of the subthalamic nucleus changes the firing pattern of pallidal neurons. J Neurosci. 2003; 23(5):1916–23.

15. Kim T, Kadji H, Whalen AJ, Ashourvan A, Freeman E, Fried SI, et al. Thermal effects on neurons during stimulation of the brain. J Neural Eng. 2022; 19(5):056029.

16. Liu Y, Postupna N, Falkenberg J, Anderson ME. High frequency deep brain stimulation: what are the therapeutic mechanisms? Neurosci Biobehav Rev. 2008; 32(3):343–51.

17. Couturier NH, Durand DM. Corpus callosum low-frequency stimulation suppresses seizures in an acute rat model of focal cortical seizures. Epilepsia. 2018; 59(12):2219–30.

18. Koubeissi MZ, Joshi S, Eid A, Emami M, Jaafar N, Syed T, et al. Low-frequency stimulation of a fiber tract in bilateral temporal lobe epilepsy. Epilepsy Behav. 2022; 130:108667.

19. Butson CR, Cooper SE, Henderson JM, McIntyre CC. Patient-specific analysis of the volume of tissue activated during deep brain stimulation. Neuroimage. 2007; 34(2):661–70.

20. McIntyre CC, Mori S, Sherman DL, Thakor NV, Vitek JL. Electric field and stimulating influence generated by deep brain stimulation of the subthalamic nucleus. Clin Neurophysiol. 2004; 115(3):589–95.

21. Dayal V, Grover T, Limousin P, Akram H, Cappon D, Candelario J, et al. The effect of short pulse width settings on the therapeutic window in subthalamic nucleus deep brain stimulation for Parkinson’s disease. J Park Dis. 2018; 8(2):273–9.

22. Elder C, Friedman D, Devinsky O, Doyle W, Dugan P. Responsive neurostimulation targeting the anterior nucleus of the thalamus in 3 patients with treatment-resistant multifocal epilepsy. Epilepsia Open. 2019; 4(1):187–92.

23. Beaudreault CP, Muh CR, Naftchi A, Spirollari E, Das A, Vazquez S, et al. Responsive Neurostimulation Targeting the Anterior, Centromedian and Pulvinar Thalamic Nuclei and the Detection of Electrographic Seizures in Pediatric and Young Adult Patients. Front Hum Neurosci. 2022; 16:876204.

24. Jin H, Li W, Dong C, Wu J, Zhao W, Zhao Z, et al. Hippocampal deep brain stimulation in nonlesional refractory mesial temporal lobe epilepsy. Seizure. 2016; 37:1–7.

25. Cukiert A, Cukiert CM, Burattini JA, Mariani PP, Bezerra DF. Seizure outcome after hippocampal deep brain stimulation in patients with refractory temporal lobe epilepsy: A prospective, controlled, randomized, double-blind study. Epilepsia. 2017; 58(10):1728–33.

26. Chen N, Zhang J-G, Han C-L, Meng F-G. Hippocampus chronic deep brain stimulation induces reversible transcript changes in a macaque model of mesial temporal lobe epilepsy. Chin Med J (Engl). 2021; 134(15):1845–54.

27. Salam MT, Velazquez JLP, Genov R. Seizure Suppression Efficacy of Closed-Loop Versus Open-Loop Deep Brain Stimulation in a Rodent Model of Epilepsy. IEEE Trans Neural Syst Rehabil Eng. 2016; 24(6):710–9.

28. Zhang F, Yang Y, Zheng Y, Zhu J, Wang P, Xu K. Combination of Matching Responsive Stimulations of Hippocampus and Subiculum for Effective Seizure Suppression in Temporal Lobe Epilepsy. Front Neurol. 2021; 12:638795.

29. Jeffrey M, Lang M, Gane J, Wu C, Burnham WM, Zhang L. A reliable method for intracranial electrode implantation and chronic electrical stimulation in the mouse brain. BMC Neurosci. 2013; 14(1):82.

30. Sobayo T, Mogul DJ. Should stimulation parameters be individualized to stop seizures: Evidence in support of this approach. Epilepsia. 2016; 57(1):131–40.

31. D’Arcangelo G, Panuccio G, Tancredi V, Avoli M. Repetitive low-frequency stimulation reduces epileptiform synchronization in limbic neuronal networks. Neurobiol Dis. 2005; 19(1–2):119–28.

32. Ruan Y, Xu C, Lan J, Nao J, Zhang S, Fan F, et al. Low-frequency Stimulation at the Subiculum is Anti-convulsant and Anti-drug-resistant in a Mouse Model of Lamotrigine-resistant Temporal Lobe Epilepsy. Neurosci Bull. 2020; 36(6):654–8.

33. Kile KB, Tian N, Durand DM. Low frequency stimulation decreases seizure activity in a mutation model of epilepsy. Epilepsia. 2010; 51(9):1745–53.

34. Alcala-Zermeno JL, Starnes K, Gregg NM, Worrell G, Lundstrom BN. Responsive neurostimulation with low-frequency stimulation. Epilepsia. 2022;:epi.17467.

35. Takeuchi Y, Harangozó M, Pedraza L, Földi T, Kozák G, Li Q, et al. Closed-loop stimulation of the medial septum terminates epileptic seizures. Brain [Internet]. 2021;. Available from: http://dx.doi.org/10.1093/brain/awaa450

36. Wang Y, Shen Y, Cai X, Yu J, Chen C, Tan B, et al. Deep brain stimulation in the medial septum attenuates temporal lobe epilepsy via entrainment of hippocampal theta rhythm. CNS Neurosci Ther. 2021; 27(5):577–86.

37. Ahn S, Jo S, Jun SB, Lee HW, Lee S. Study on the mechanisms of seizure-like events suppression effect by electrical stimulation using a microelectrode array. Neuroreport. 2017; 28(9):471–8.

38. Boёx C, Vulliémoz S, Spinelli L, Pollo C, Seeck M. High and low frequency electrical stimulation in non-lesional temporal lobe epilepsy. Seizure. 2007; 16(8):664–9.

39. Donos C, M\^\indru\c tă I, Ciurea J, Măl\^\iia MD, Barborica A. A comparative study of the effects of pulse parameters for intracranial direct electrical stimulation in epilepsy. Clin Neurophysiol. 2016; 127(1):91–101.

40. Basu I, Robertson MM, Crocker B, Peled N, Farnes K, Vallejo-Lopez DI, et al. Consistent linear and non-linear responses to invasive electrical brain stimulation across individuals and primate species with implanted electrodes. Brain Stimul. 2019; 12(4):877–92.

41. Cogan SF, Ludwig KA, Welle CG, Takmakov P. Tissue damage thresholds during therapeutic electrical stimulation. J Neural Eng. 2016; 13(2):021001.

42. Quiroga RQ, Nadasdy Z, Ben-Shaul Y. Unsupervised Spike Detection and Sorting with Wavelets and Superparamagnetic Clustering. Neural Comput. 2004; 16(8):1661–87.

43. Pfammatter JA, Bergstrom RA, Wallace EP, Maganti RK, Jones MV. A predictive epilepsy index based on probabilistic classification of interictal spike waveforms. Biagini G, editor. PLOS ONE. 2018; 13(11):e0207158.

44. Cimbalnik J, Klimes P, Sladky V, Nejedly P, Jurak P, Pail M, et al. Multi-feature localization of epileptic foci from interictal, intracranial EEG. Clin Neurophysiol. 2019; 130(10):1945–53.

45. Huneau C, Benquet P, Dieuset G, Biraben A, Martin B, Wendling F. Shape features of epileptic spikes are a marker of epileptogenesis in mice. Epilepsia. 2013; 54(12):2219–27.

46. Seabold S, Perktold J. Statsmodels: Econometric and Statistical Modeling with Python. {Millman”] [“Stéfan van der Walt, Jarrod}, editors. 2010;:92–6.

47. Monaghan DT, Cotman CW. The distribution of [3H]kainic acid binding sites in rat CNS as determined by autoradiography. Brain Res. 1982; 252(1):91–100.

48. McIntyre CC, Savasta M, Kerkerian-Le Goff L, Vitek JL. Uncovering the mechanism(s) of action of deep brain stimulation: activation, inhibition, or both. Clin Neurophysiol. 2004; 115(6):1239–48.

49. Toprani S, Durand DM. Fiber tract stimulation can reduce epileptiform activity in an in-vitro bilateral hippocampal slice preparation. Exp Neurol. 2013; 240:28–43.

50. Paxinos G, Franklin K. The Mouse Brain in Stereotaxic Coordinates. 2nd ed. Academic Press, San Diego.; 2001.

